# Consensus Co-Expression Analysis Identifies A Common Set Of Co-Expressed Genes Associated With Diabetic Peripheral Neuropathy And Chemotherapy-Induced Peripheral Neuropathy

**DOI:** 10.1101/2025.11.18.689101

**Authors:** Kord M. Kober, Esther Chavez-Iglesias, Nam Woo Cho, Sue Yom, Niharika Dixit, Marina Sirota, Alexandre Chan, Adam Olshen

**Author notes:** Address correspondence to: Kord M. Kober, Associate Professor, 521 Parnassus Avenue, 3229, Box 0638, San Francisco, California 94117.

## Abstract

**Background:** Diabetic peripheral neuropathy (DPN) and chemotherapy-induced peripheral neuropathy (CIPN) are major clinical challenges with limited therapeutic options. While these conditions arise from different causes, they may share common molecular mechanisms that could be targeted for intervention.

**Methods:** We performed consensus weighted gene co-expression network analysis (WGCNA) on two publicly available datasets: GSE185011 (DPN vs. healthy controls in peripheral blood mononuclear cells) and GSE173610 (paclitaxel-treated vs. control iPSC-derived sensory neurons). After filtering all but the most variable genes, consensus analysis was used to identify conserved co-expression modules across both conditions.

**Results:** Consensus analysis identified a 193-gene module (ME3/brown) significantly associated with both DPN (correlation=0.817, p=0.0040) and CIPN (correlation=0.971, p=0.0060). Functional enrichment analysis of this module revealed pathways related to Glycolysis, FoxO signaling, Apoptosis, and Autophagy.

**Conclusions:** Our analysis reveals a convergent molecular signature underlying both DPN and CIPN, centered on metabolic reprogramming, transcriptional stress, and programmed cell death. These findings provide a systems-level framework for developing therapies targeting shared pathological mechanisms.

## Introduction

Peripheral neuropathy affects millions of patients worldwide, with diabetic peripheral neuropathy (DPN) and chemotherapy-induced peripheral neuropathy (CIPN) representing two of the most prevalent forms. DPN occurs in approximately 50% of diabetic patients and is characterized by progressive sensory loss, neuropathic pain, and increased risk of foot ulceration and amputation.^1-3^ CIPN affects 30-70% of patients receiving neurotoxic chemotherapeutic agents such as taxanes, platinum compounds, and vinca alkaloids, often leading to dose reductions or treatment discontinuation that may compromise cancer outcomes.^4, 5^

Despite their different etiologies (i.e., chronic metabolic dysfunction in DPN^6^ versus acute neurotoxic injury in CIPN^7^) both conditions share remarkably similar clinical presentations, including distal sensory loss, neuropathic pain, and a characteristic “stocking-glove” distribution of symptoms. This phenotypic convergence suggests that these distinct insults may trigger common downstream molecular pathways leading to neuronal dysfunction and death.

Current therapeutic approaches for both conditions remain largely symptomatic, focusing on pain management rather than addressing underlying pathophysiology.^6, 8-12^ The lack of disease-modifying treatments partly reflects our incomplete understanding of the molecular mechanisms driving neuronal injury in these conditions.^13^ While individual studies have identified specific pathways involved in DPN^14^ or CIPN,^15-17^ a systematic comparison of their molecular signatures has not been performed.

Modern systems biology approaches, particularly weighted gene co-expression network analysis (WGCNA),^18, 19^ provide powerful tools for identifying conserved biological modules across different disease states. Unlike traditional differential expression analysis, WGCNA identifies clusters of co-expressed genes that function together, providing insights into coordinated biological processes and regulatory networks.

The objective of this study was to identify shared molecular mechanisms between DPN and CIPN using consensus co-expression analysis of publicly available transcriptomic datasets. We hypothesized that despite their different etiologies, these neuropathies would converge on stress pathways that could represent universal therapeutic targets for neuroprotection.

## Materials and Methods

### Data Acquisition and Preprocessing

Transcriptomic data from two publicly available peripheral neuropathy studies consisted of RNA-sequencing data obtained from the Gene Expression Omnibus (GEO) database.^20^ The first dataset, GSE185011, profiled gene expression in peripheral blood mononuclear cells from patients with DPN and healthy controls.^14^ The second dataset, GSE173610, from human induced pluripotent stem cell (iPSC)-derived sensory neurons treated with paclitaxel to induce CIPN, with DMSO-treated cells serving as controls.^21, 22^

For GSE185011, raw FPKM values were provided with gene symbols aggregated by taking the mean of multiple entries. For GSE173610, TPM values were provided and mapped from Ensembl IDs to gene symbols using org.Hs.eg.db, retaining the ID with highest expression variance when multiple IDs mapped to the same symbol. Both datasets were independently filtered to retain the top 7,000 genes with highest variance across samples.

### Individual and Consensus WGCNA

Traditional pathway-based analyses are constrained by existing biological knowledge and may miss novel regulatory networks. WGCNA addresses this limitation by examining genome-wide expression patterns to detect clusters of genes that exhibit coordinated expression behaviors across samples. These gene clusters, termed modules, are identified through unsupervised clustering methods that group genes based on expression similarity rather than predetermined biological categories. The resulting co-expression modules often reflect shared regulatory mechanisms or functional relationships, and when correlated with phenotypic traits, can reveal previously unrecognized molecular signatures underlying these adverse conditions. Individual WGCNA was performed on each preprocessed dataset using the WGCNA R package. ^18, 19^ After sample clustering to identify outliers, soft-thresholding powers were selected based on scale-free topology model fit. Adjacency matrices were calculated and converted to topological overlap matrices (TOM). Hierarchical clustering with the Dynamic Tree Cut method^23^ (minimum module size = 30 genes) identified gene modules, with highly correlated modules merged (cut height = 0.25).

Consensus WGCNA was performed on the intersection of genes present in both datasets using blockwise consensus module detection. For each dataset, consensus module eigengenes were calculated and correlated with the respective clinical trait. A “convergent module” was defined as any module showing a statistically significant (p < 0.05) correlation with the clinical trait in both datasets. Among these, a single “key” module was selected for downstream analysis by identifying the module with the highest geometric mean of its absolute trait correlations across all datasets. Consensus hub genes for each convergent module were identified based on their module membership, or kME (Module Eigengene connectivity), calculated across both datasets using the consensusKME function. All genes were ranked in the module based on the absolute value of the single, consensus kME score and the top 30 genes from this consensus ranking were selected as the hub genes.

### Functional Analysis

Protein-protein interaction networks were constructed using STRING DB (version 12.0, medium confidence > 0.400).^24^ Pathway enrichment analysis was performed using KEGG,^25^ Reactome,^26^ WikiPathways^27^ databases, and Gene Ontology (GO) terms.^28^

## Results

A total of 2,630 genes overlapped between datasets and were included in consensus analysis. Two modules showed significant associations with neuropathy traits: ME3/brown (geometric mean of absolute correlations: 0.981, correlation=0.971, p=0.0060 for CIPN; correlation=0.817, p=0.0040 for DPN) and ME26/darkorange (geometric mean of absolute correlations: 0.800, correlation=0.949, p=0.0136 for CIPN; correlation=0.674, p=0.0325 for DPN). The ME3/brown module was selected as the key consensus module, containing 193 genes. Key hub genes (Supplemental File 2) included GAPDH, PKM2, HK1, ENO1, FOXO3, CREB1, KDM6B, KMT2C, SRSF2, SF3B1, SARM1, ULK1, BBC2 (PUMA).

The module-trait correlation heatmap highlighting the convergent brown module is shown in Figure 1A. Figure 1B demonstrates consistent elevation of the brown module’s eigengene expression in both neuropathy conditions compared to controls. Network preservation statistics (Figure 1C) confirm robust preservation across datasets. Figure 1D displays the network of the genes within the brown module.

**Figure 1.**
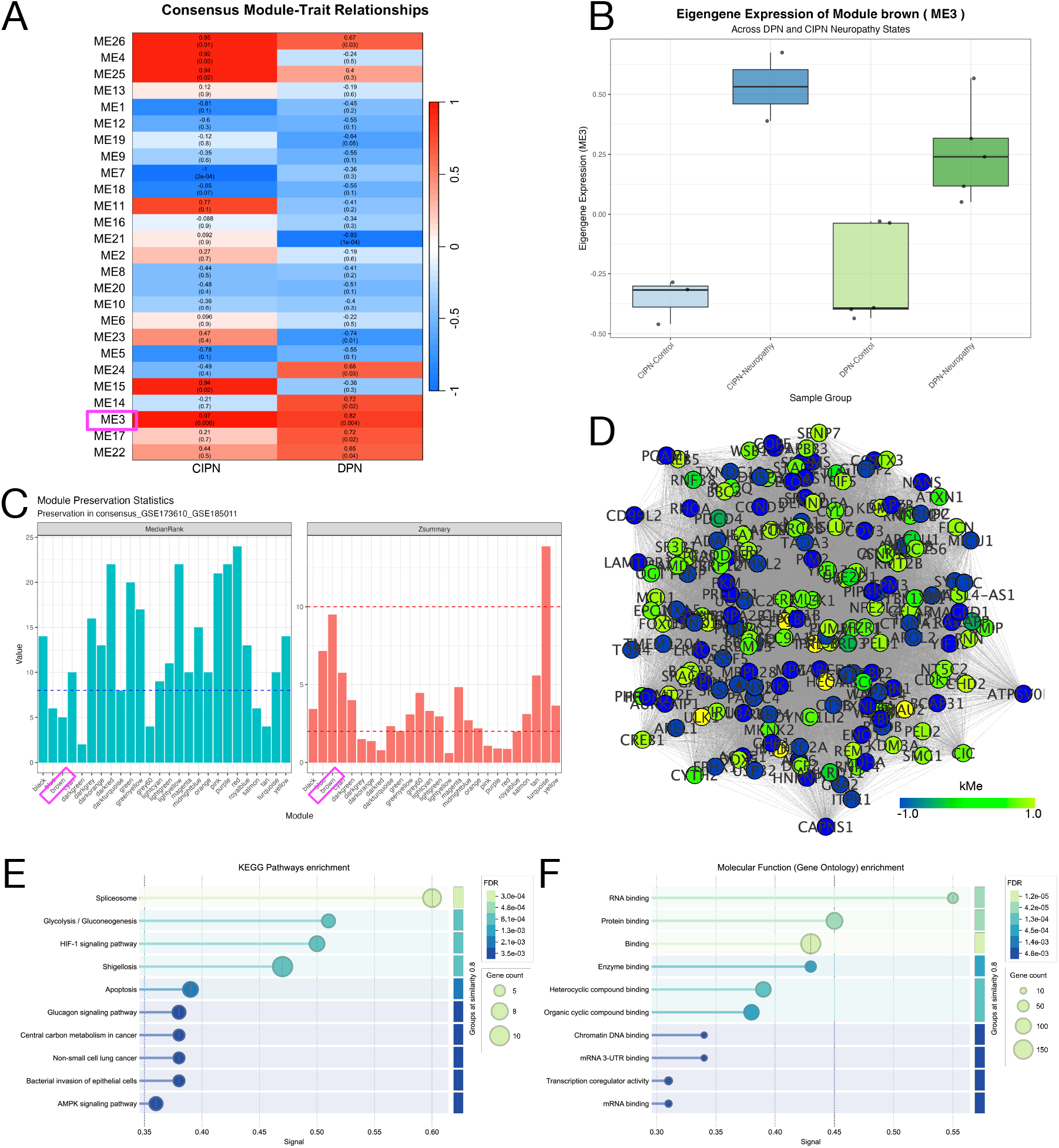
Consensus Co-Expression Module identified across chemotherapy-induced and diabetic peripheral neuropathies. (A) Module-trait heatmap across the two studies. The key consensus module is marked with a magenta square. (B) Module eigengene expression across the two studies. (C) Network preservation statistics. (D) Network visualization of the consensus module. Edge widths are scaled by TOM weights. Nodes are colored by eigengene-based (kME) value. (E) Top 10 enriched KEGG pathways for genes in the consensus module. (F) Top 10 enriched GO Molecular Function terms for genes in the consensus module.

Beyond the module-trait associations, the protein-protein interaction network was significantly enriched (p < 1.0×10^−16^) with 191 nodes, 411 edges, and an average clustering coefficient of 0.399 (Supplemental Table 1, Supplemental Figure 1). Functional enrichment analysis of the brown module identified significant enrichments for 104 Biological Processes, 12 Molecular Functions, and 29 Cellular Component GO terms (Supplemental Figure 2, Supplemental File 3), as well as 14 KEGG, 4 Reactome, and 10 WikiPathways (Supplemental Figure 3, Supplemental File 3).

## Discussion

This study represents the first systematic identification of shared molecular mechanisms between diabetic peripheral neuropathy and chemotherapy-induced peripheral neuropathy through consensus co-expression network analysis. Despite their distinct etiologies—chronic metabolic dysregulation in DPN versus acute cytotoxic injury in CIPN—these conditions manifest with a remarkably similar clinical phenotype of length-dependent, sensory-predominant axonopathy,^29^ often characterized by numbness, tingling, burning pain, and a “glove-and-stocking” distribution.^30^ Our analysis provides a systems-level explanation for this clinical convergence by identifying a robustly preserved 193-gene co-expression module whose activity is significantly associated with both disease states.

### A Convergent Pathophysiological Framework: From Metabolic Crisis to Neurodegeneration

The functional architecture of the brown module reveals three interconnected pathological themes: metabolic reprogramming centered on glycolysis, systemic dysregulation of gene expression driven by FoxO signaling, and activation of programmed cell death pathways including apoptosis and autophagy. This framework suggests that while initial triggers differ—chronic hyperglycemia and dyslipidemia in DPN versus direct mitochondrial damage from cytotoxic agents in CIPN—they converge on a common downstream pathway involving oxidative stress, mitochondrial dysfunction, and cellular demise.^1, 6^

### Metabolic Crisis and Bioenergetic Failure

The enrichment for glycolytic pathways (e.g., “Glycolysis / Gluconeogenesis” and “Glycolysis”), including key enzymes glyceraldehyde-3-phosphate dehydrogenase (GAPDH), pyruvate kinase M2 (PKM2), hexokinase 1 (HK1), and enolase 1 (ENO1), underscores fundamental disruption of cellular energy homeostasis as a primary convergence point. The coordinated upregulation of glycolytic genes likely represents cellular compensation for mitochondrial dysfunction, particularly noted in Schwann cells supporting injured axons.^31, 32^

However, these enzymes extend beyond canonical metabolic roles through critical “moonlighting” functions as signaling molecules and mediators of cell death.^33^ Under hyperglycemic conditions, mitochondrial superoxide overproduction inhibits GAPDH, diverting upstream metabolites into damaging pathways, including advanced glycation end product formation.^34^ Furthermore, under nitrosative stress, S-nitrosylated GAPDH translocates to the nucleus to initiate cell death cascades.^34, 35^ GAPDH’s central position within the convergent network suggests its pathological modification represents a critical node where distinct insults trigger neurodegeneration.^36, 37^

### Transcriptional Dysregulation and Proteostasis Disruption

The neuropathic state is characterized by system-wide transcriptional reprogramming, evidenced by enrichment for FoxO signaling pathways and FOXO-mediated transcription.^38^ This regulation involves master transcription factors, such as FOXO3 and CREB1, as well as chromatin-modifying enzymes including KDM6B and KMT2C, and RNA processing machinery components, including splicing factors SRSF2 and SF3B1. The coordinated dysregulation suggests maladaptive rewiring of cellular informational architecture, ^39^ where both chronic metabolic stress and acute cytotoxic insults overwhelm precise genomic responses.^40, 41^

This maladaptive rewiring may lead to transcriptional and post-transcriptional dysregulation, including aberrant RNA splicing (e.g., “mRNA Splicing – Major Pathway” and “Spliceosome”), which generates dysfunctional proteins and contributes to cellular pathology.^42, 43^ The presence of both regulatory “command” elements and “execution” machinery within the same co-expression module highlights heightened yet potentially dysfunctional transcriptional activity contributing to progressive neurodegeneration.^44^ Emerging evidence suggests that alternative splicing plays a role in major neurodegenerative diseases, indicating that this mechanism may be a promising target for developing therapies.^43^

### Programmed Cell Death Pathway Activation

The ultimate downstream consequence involves activation of programmed cell death pathways (e.g., “Apoptosis,” “FOXO-mediated transcription of cell death genes,” and “Autophagy - animal”). ^6, 45^ Specific genes within the module that highlight convergence on core cellular self-destruction machinery include the pro-apoptotic BH3-only protein BBC3, anti-apoptotic MCL1, and autophagy initiator ULK1.^46^ The co-expression of these regulators suggests the brown module represents a critical decision-making hub integrating opposing signals and determining cell fate. In DPN and CIPN, persistent overwhelming stress shifts this balance toward pro-death signaling and progressive neurodegeneration.^47, 48^

### Key Hub Genes as Therapeutic Targets

Several highly connected hub genes (i.e., FOXO3, PKM2, HK1, ULK1, BBC3) emerge as critical nodes integrating pathological themes and representing therapeutic opportunities. First, FOXO3 serves as the master regulator connecting all three themes: sensing metabolic crisis,^49, 50^ executing transcriptional dysregulation,^51, 52^ and triggering programmed cell death by transcriptionally upregulating pro-apoptotic BH3-only proteins, including BBC3 (also known as the p53 upregulated modulator of apoptosis (PUMA)).^52, 53^ Next, the glycolytic enzymes PKM2 (pyruvate kinase M2) and HK1 (hexokinase 1), which are critical for glucose metabolism, are highly connected hub genes, suggesting their centrality to convergent pathology. Their potential role lies not only in canonical energy production but in non-canonical “moonlighting” roles as active mediators of cell signaling and death.^33, 54^ PKM2 represents a metabolic hub with non-canonical signaling roles. Beyond glycolytic functions, dimeric PKM2 translocates to the nucleus as a transcriptional co-activator influencing inflammation-related gene expression. ^55, 56^ PKM2 is implicated in inflammatory processes driving central sensitization in neuropathic pain models.^57-59^ Hexokinases (HKs) catalyze the first step of glucose metabolism, whereby glucose enters glycolysis via phosphorylation by HK1.^60^ HK1 provides direct genetic evidence linking metabolic dysregulation to neuropathy, as mutations cause Charcot-Marie-Tooth disease with phenotypes similar to DPN and CIPN. ^61^

Finally, ULK1 and BBC3 represent gatekeepers of cellular degradation programs. ULK1 initiates autophagy for neuronal quality control ^62, 63^ but also regulates SARM1-mediated axonal degeneration.^64, 65^ BBC3 is a powerful direct activator of intrinsic mitochondrial apoptosis^66, 67^ transcriptionally upregulated by FOXO3.^53^ The simultaneous presence of ULK1, the primary autophagy initiator, and BBC3, a potent apoptosis initiator, within the brown module points to coordinated activation of cellular demolition programs.

### Therapeutic Implications and Universal Neuroprotective Strategies

The identification of a convergent molecular network driven by specific, druggable hub genes provides a compelling rationale for a paradigm shift in the treatment of peripheral neuropathy. Rather than pursuing cause-specific strategies—such as glycemic control in DPN or mitigating microtubule disruption in CIPN—a more effective approach may be targeting downstream molecular nodes where distinct pathologies converge.^68, 69^ Our analysis highlights several promising targets for universal, etiology-independent neuroprotective therapies.^69^ Given the lack of effective established agents for preventing or treating DPN and CIPN, and the increased prevalence of diabetes ^70^ and the increasing number of cancer survivors, there is an urgent need for the identification and development of new, effective intervention strategies.^11, 71^

### Study Limitations

Several limitations should be acknowledged. Our analysis relied on different tissue types (peripheral blood mononuclear cells for DPN versus iPSC-derived sensory neurons for CIPN), which may have introduced bias toward systemically expressed genes. Small sample sizes limit statistical power and generalizability. The analysis is correlative and does not establish causality. Future studies should validate findings in independent cohorts using consistent tissue types such as dorsal root ganglia. Functional validation through targeted manipulation in relevant models will establish causality and therapeutic potential. Integration with proteomic and metabolomic data would provide more comprehensive molecular views.

### Clinical Translation Potential

Rather than developing separate therapeutic approaches for each neuropathy type, our findings suggest universal neuroprotective strategies targeting shared pathways could benefit patients regardless of etiology. This approach could be particularly valuable for patients with multiple risk factors or neuropathy of unclear cause. The prominence of stress response pathways aligns with emerging therapeutic strategies focused on enhancing cellular resilience rather than targeting specific disease mechanisms. Approaches such as hormesis, preconditioning, and pharmacological chaperone therapy could provide broad neuroprotective benefits by strengthening intrinsic stress response capabilities.

## Conclusions

This study establishes a robust framework for identifying conserved molecular mechanisms across different neuropathy etiologies and demonstrates that DPN and CIPN converge on common pathways involving metabolic reprogramming, transcriptional stress, and programmed cell death. The identification of druggable targets within this convergent network provides a foundation for developing universal neuroprotective therapies that could transform the treatment landscape for peripheral neuropathy. By shifting from cause-specific to mechanism-based therapeutic approaches, these findings open new avenues for developing treatments that could benefit millions of patients suffering from these debilitating conditions.

## Acknowledgments

Research reported in this study was supported by grants from the National Cancer Institute (R37, CA233774) and a Cancer Center Support Grant (P30, CA082103). Its contents are solely the responsibility of the authors and do not necessarily represent the official views of the NIH.

The async development agent Jules (http://jules.google.com) was used to support software development, debugging, testing, generating tables and figures, and documenting the project. Google Gemini (http://gemini.google.com, LLM version 2.5 Pro) was utilized to brainstorm initial study designs and refine the summarization of findings, under direct human supervision. Anthropic General-Purpose Assistant (LLM version Claude Sonnet 4) via UCSF Versa was utilized to assist in the language of the manuscript. Jenni.AI (https://jenni.ai/) was used to assist in the literature search.

## Author Contributions

*KMK: Conceptualization, Data curation, Data curation, Funding acquisition, Investigation, Methodology, Project administration, Resources, Software, Supervision, Validation, Visualization, Writing – original draft, Writing – review & editing*

*ECI: review & editing, Validation*

*NWC: Writing – review & editing, Validation*

*SY: Writing – review & editing, Validation*

*ND: Writing – review & editing, Validation*

*MS: Writing – review & editing, Validation*

*AC: Writing – review & editing, Validation*

*AO: Formal analysis, Methodology, Writing – review & editing, Validation, Resources*

## Disclosures

The author(s) declare no potential conflicts of interest with respect to the research, authorship, and/or publication of this article.

## Notes

### Competing Interest Statement

The authors have declared no competing interest.

