## Supplemental File 1 for "Consensus Co-Expression Analysis Identifies A Common Set Of Co-Expressed Genes Associated With Diabetic Peripheral Neuropathy And Chemotherapy-Induced Peripheral Neuropathy"

**This PDF file includes:**

Supplemental Table 1. Network summary statistics for the STRING DB protein-protein interaction network of the Brown consensus module genes
Supplemental Figure 1. Protein-protein interaction network for key Brown consensus module
Supplemental Figure 2. Top 10 GO enriched terms for key Brown consensus module
Supplemental Figure 3. Top 10 KEGG, Reactome, WikiPathway enriched pathways for key Brown consensus module

**Supplemental Data Tables (in separate documents):**

Supplemental File 2. List of genes in the Brown module.
Supplemental File 3. Enrichment of GO terms and KEGG, Reactome, and WikiPathways pathways for genes in the Brown consensus module.

Supplemental Table 1. Network summary statistics for the STRING DB protein-protein interaction network of the consensus module genes.

| Characteristic | Score |
|---|---|
| Number of nodes | 160 |
| Number of edges | 411 |
| Avg. number of neighbors | 5.423 |
| Network diameter | 8 |
| Network radius | 4 |
| Characteristic path length | 3.488 |
| Clustering coefficient | 0.249 |
| Network density | 0.037 |
| Network heterogeneity | 0.957 |
| Network centralization | 0.223 |
| Connected components | 5 |

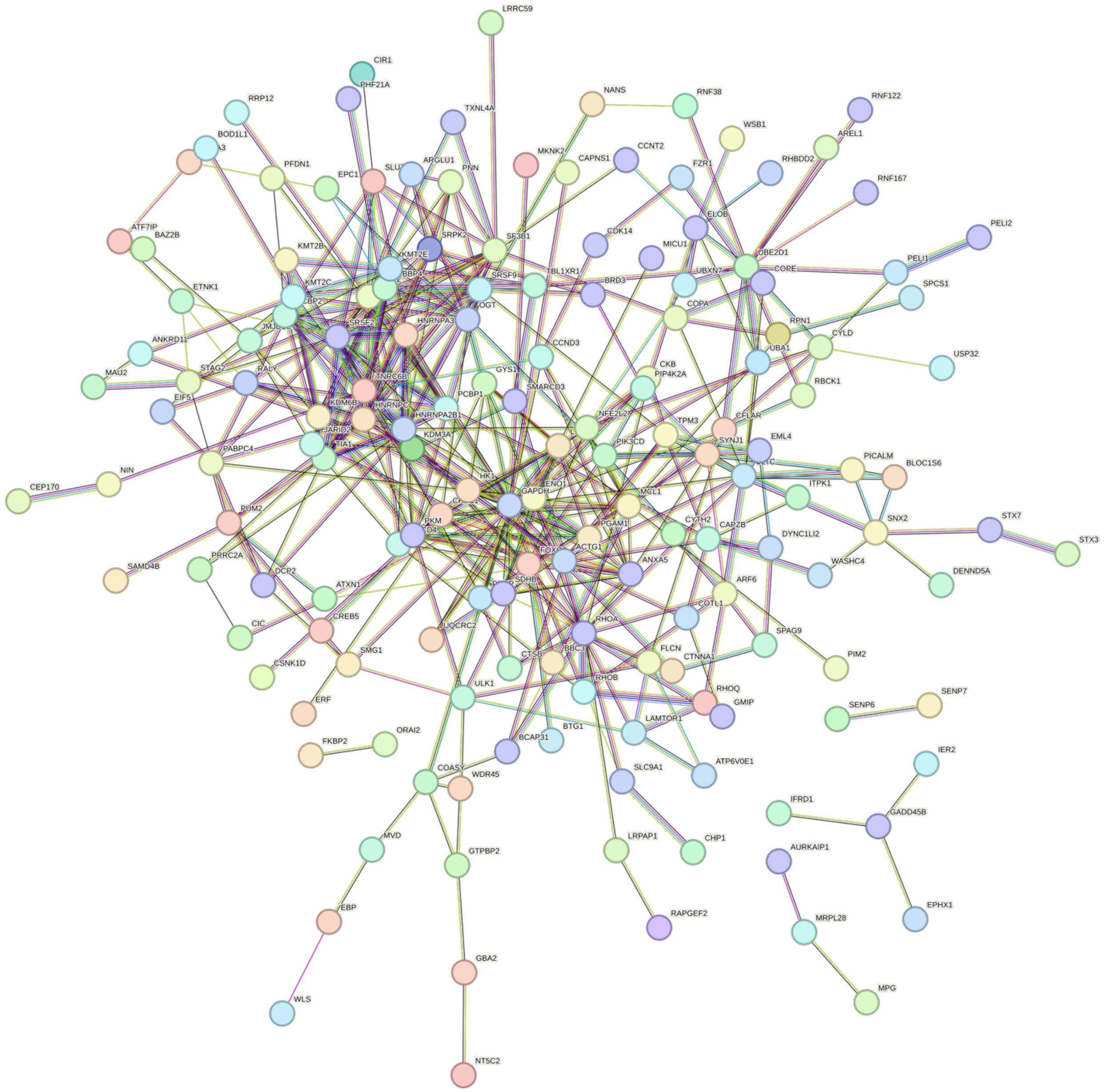


Supplemental Figure 1. Protein-protein interaction network for genes in the Brown consensus module across diabetic peripheral neuropathy and chemotherapy-induced peripheral neuropathy.

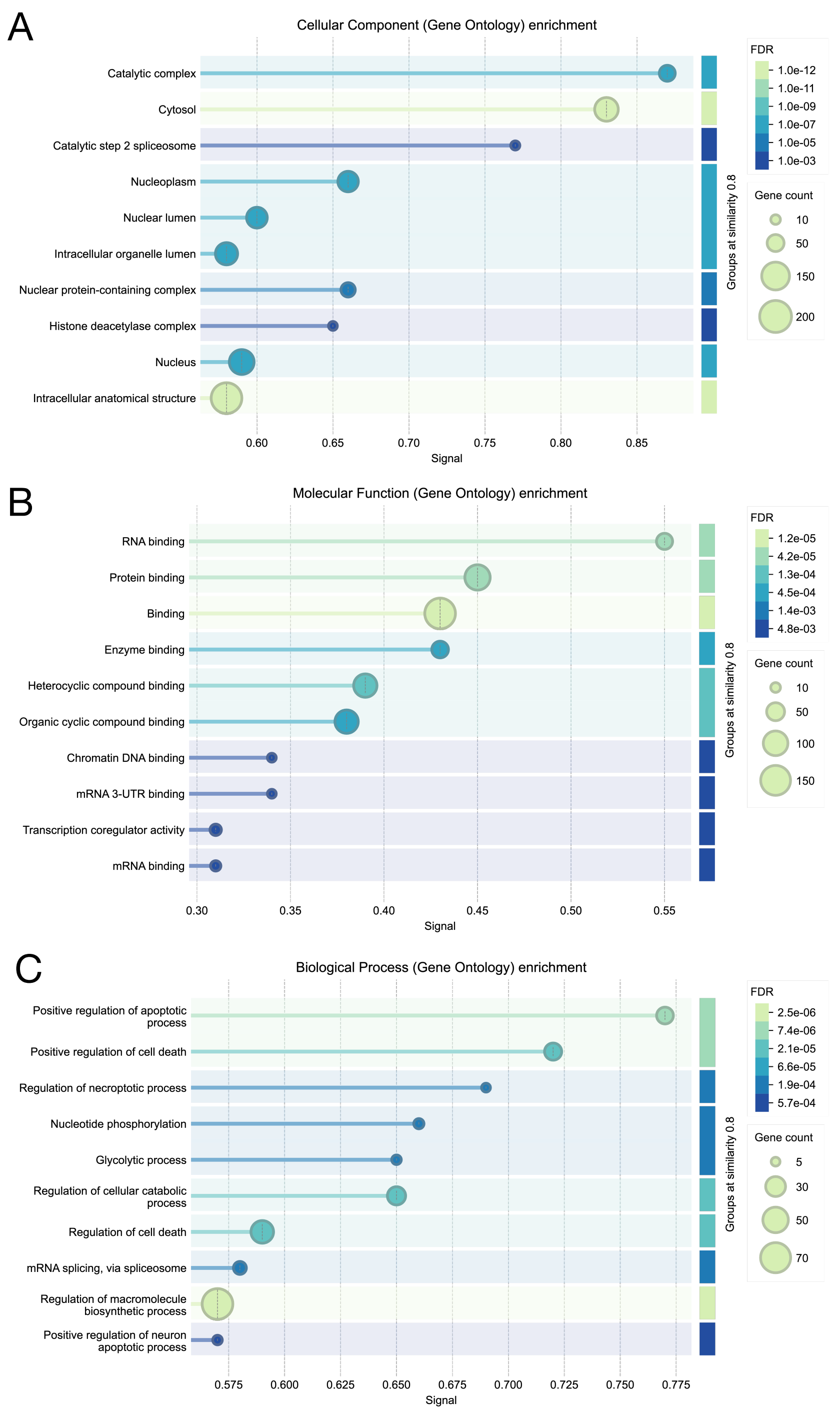


Supplemental Figure 2. Enrichment of GO terms in Molecular Function, Cellular Component, and Biological Processes for genes in the Brown consensus module across diabetic peripheral neuropathy and chemotherapy-induced peripheral neuropathy.

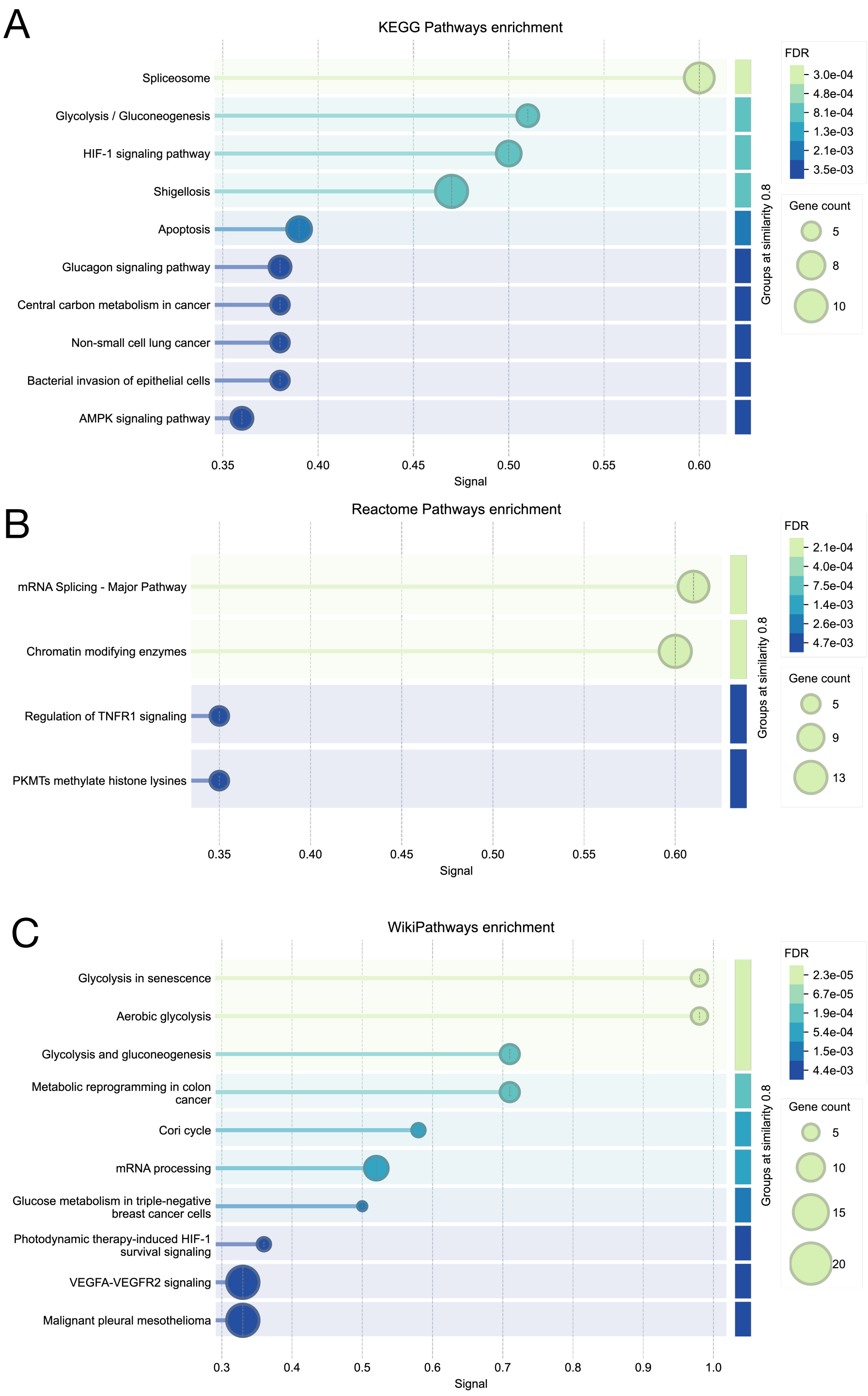


Supplemental Figure 3. Enrichment of KEGG, Reactome, and WikiPathway database pathways for genes in the Brown consensus module across diabetic peripheral neuropathy and chemotherapy-induced peripheral neuropathy.
